# Subcellular mapping of the protein landscape of SARS-CoV-2 infected cells for target-centric drug repurposing

**DOI:** 10.1101/2022.03.29.482838

**Authors:** Jayasankar Mohanakrishnan Kaimal, Marianna Tampere, Trang H. Le, Ulrika Axelsson, Hao Xu, Hanna Axelsson, Anna Bäckström, Francesco Marabita, Elisabeth Moussaud-Lamodière, Duncan Njenda, Carolina Oses Sepulveda, Wei Oyuang, Brinton Seashore-Ludlow, Caroline Vernersson, Ali Mirazimi, Emma Lundberg, Päivi Östling, Charlotte Stadler

## Abstract

The COVID-19 pandemic has resulted in millions of deaths and affected socioeconomic structure worldwide and the search for new antivirals and treatments are still ongoing. In the search for new drug target and to increase our understanding of the disease, we used large scale immunofluorescence to explore the host cell response to SARS-CoV-2 infection. Among the 602 host proteins studied in this host response screen, changes in abundance and subcellular localization were observed for 97 proteins, with 45 proteins showing increased abundance and 10 reduced abundances. 20 proteins displayed changed localization upon infection and an additional 22 proteins displayed altered abundance and localization, together contributing to diverse reshuffling of the host cell protein landscape. We then selected existing and approved small-molecule drugs (n =123) against our identified host response proteins and identified 3 compounds - elesclomol, crizotinib and rimcazole, that significantly reduced antiviral activity. Our study introduces a novel, targeted and systematic approach based on host protein profiling, to identify new targets for drug repurposing. The dataset of ∼75,000 immunofluorescence images from this study are published as a resource available for further studies.

## Introduction

The severe acute respiratory syndrome coronavirus-2 (SARS-CoV-2) causing COVID-19 has led to more than 5 million deaths (https://covid19.who.int/) and initiated an unforeseen health and socioeconomic crisis (1). Such continued threats by emerging pathogens emphasize the need for new approaches to comprehensively identify therapeutic targets and drug candidates.

Upon infection, viruses commonly hijack host cell machinery to enable replication. This leads to re-organization of the cellular proteome composition (2,3) such as up- or downregulation of signaling pathways (4–6) which can be quantified by large-scale omics methods. Additionally, host cells may alter abundance of proteins related to cellular defense and homeostasis. Accumulating evidence highlights the importance of host protein translocations from one organelle to another during viral infection, contributing either to host protection or viral replication (7). While bulk omics analyses capture the systematic changes, they lack spatial resolution at a single cell level and thereby information about infection-induced phenotypic changes of cellular components and translocations of specific host proteins within the cell. This information can provide deeper insight into the host cell response to viral infection since protein location is often correlated with function (8–11). Employing systematic *in situ* methodologies for studying virus-host interactions can also reveal host protein targets required by the virus for further replication and thus have implications for antiviral drug target identification, as well as drug repurposing and discovery efforts.

Publications during the first year of the pandemic have reported hundreds of host cell proteins directly interacting with at least one of the 31 SARS-CoV-2 viral proteins (12–18). These interactions were mainly studied by affinity capture methods and protein tagging of bait proteins.

In this study we investigated host cell responses upon infection using immunofluorescence and antibodies from the Human Protein Atlas (HPA)(8,9,19,20) to map the changes of host protein abundance levels and subcellular localization upon infection with SARS-CoV-2.

Additionally, we selected existing and approved small-molecule drugs against our identified altered host proteins and identified compounds with antiviral activity.

Our study introduces a novel systematic approach based on spatial protein profiling to identify novel host targets for drug repurposing, here demonstrated for SARS-CoV-2.

## Results

### SARS-CoV-2 infection affects diverse cellular functions and pathways

In order to better understand the interplay between SARS-CoV-2 and the host cell machinery, we developed an image-based assay to detect SARS-CoV-2 infection in Vero E6 cells, followed by high-resolution immunofluorescence microscopy to investigate changes in the subcellular localization and abundance of host cell proteins upon infection **(Figure 1a)**. The host cell proteins were selected and included in the host-response screen based on literature mining of previous reports identifying cellular proteins interacting with SARS-CoV-2 proteins (14–18). Building on the unique antibody resources generated within the HPA project (www.proteinatlas.org) and our established workflow for systematic subcellular mapping of proteins (8,19), 602 antibodies targeting proteins encoded by 662 genes (25 multi-targeting for highly similar proteins within the same family) were immunostained in mixed populations of infected and non-infected Vero E6 cells 24 hours after introduction of SARS-CoV-2 to the cell culture (**Supplementary table 1**). For the multi-targeting antibodies, at least one of the targets had previously been identified to interact with SARS-CoV-2. After confocal microscopy, images were uploaded into an in-house developed Covid Image Annotator tool in the ImJoy platform (21). Using a DPNUnet model, individual cells were segmented and labeled as infected and non-infected based on staining intensity of the SARS-CoV-2 nucleocapsid protein. Host protein staining intensities were then quantified and compared between infected and non-infected cells to identify altered protein abundance. Staining patterns were manually annotated to assess changes in subcellular location between the populations (**Figure 1a, Supplementary Figure 1**). The complete image dataset of the host response screen is available at *Figshare* data portal (22).

**Figure 1.**
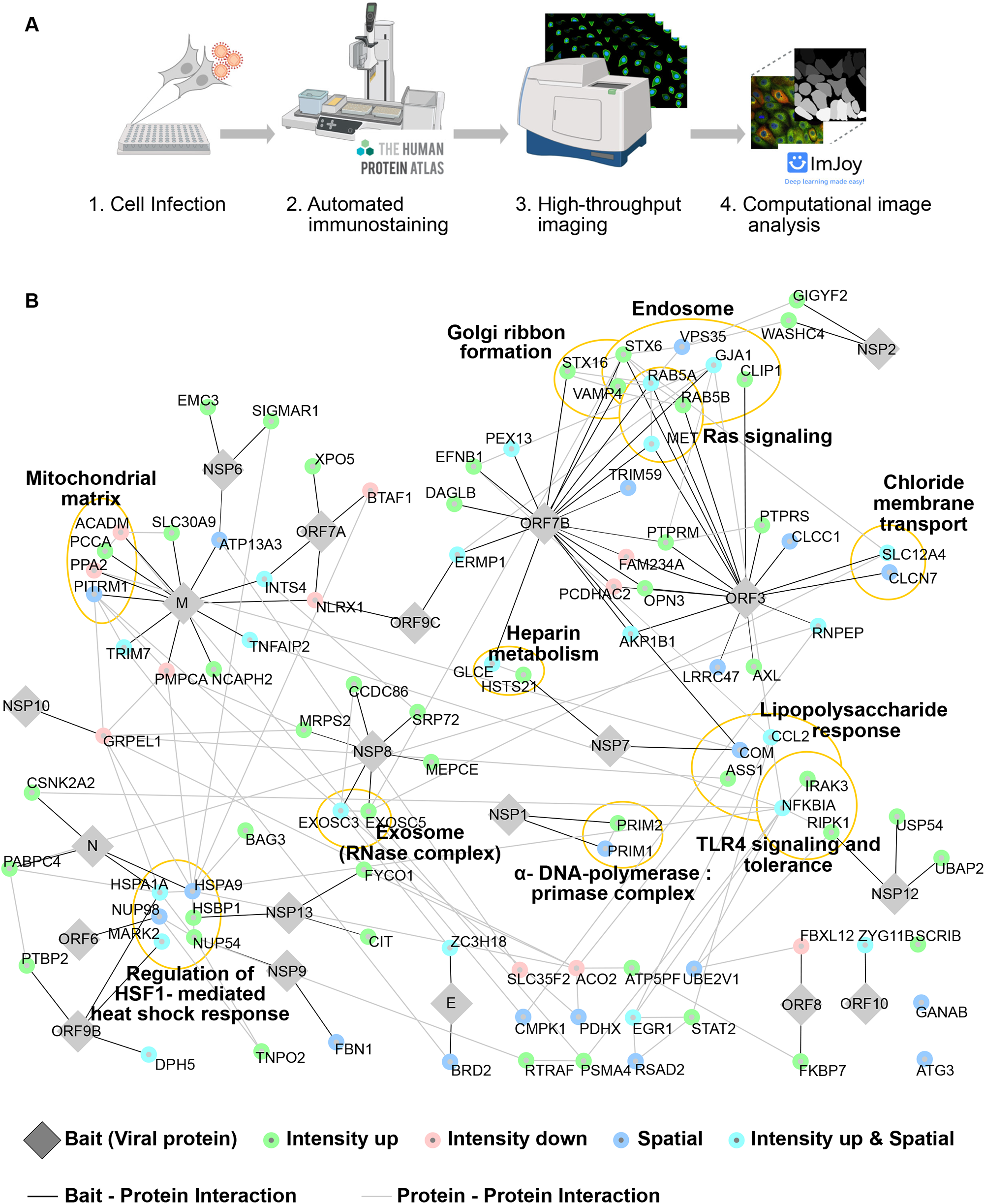
Immunofluorescence-based screen confirms complex interaction networks between SARS-CoV-2 and host proteins. A) Overview of the workflow starting from SARS-CoV-2 infection in Vero E6 cell line followed by downstream immunostaining, imaging and computational analysis. B) Protein-protein interaction network showing the 97 proteins that were altered upon infection with SARS-CoV-2. Protein clusters belonging to different GO terms (biological process, cellular components, molecular function, KEGG pathway and reactome pathway) are highlighted.

By mapping the host response of 602 proteins, we identified 97 proteins with either changed subcellular location (spatial redistribution) or altered abundance (defined as a significant difference in staining intensity) between the infected and non-infected cells. By combining our observations for the 97 proteins with the results on host cell protein interactions from literature with specific viral proteins ^18^ we generated a network map of the host cell-virus interactions using Cytoscape(23) (**Figure 1B)**.

Among the 97 proteins, most connections are linked to the SARS-CoV-2 components ORF3, ORF7B and membrane (M) protein. The M protein is one of the structural components common to all coronaviruses. ORF3 and ORF7B belong to the accessory factors, which tend to be non-essential for viral replication, but important for pathogenesis and virus-host interactions (24,25), and both ORF3 and ORF7B have been shown to modulate host cell immune responses (26,27). The host responses to SARS-CoV-2 are associated with multiple cellular components and diverse cellular functions (**Figure 1B**). Grouping and functional enrichment analysis of responding proteins includes proteins localizing to endosomes and mitochondria, and for factors involved in Golgi ribbon formation, Ras signaling, TLR4 signaling, heat shock response, lipopolysaccharide response, heparin metabolism and chloride membrane transport according to the Gene Ontology (GO), Reactome, and KEGG databases. This is in agreement with the known ability of coronaviruses to manipulate the host cell at various levels(28,29).

For example, multiple host proteins corresponding to the following genes STX6, VAMP4, VPS35, RAB5A, RAB5B, CLIP1 and GJA1 that are known to be involved in endosomal functions, displayed increased abundance as well as subcellular re-location following SARS-CoV-2 infection (**Supplementary Table 1, Figure 1B)**. This observation is in agreement that endosome formation is known to be important for host cell entry and systemic infection by coronaviruses (30,31). Furthermore, SARS-CoV-2 infection resulted in increased abundance of the proteins from the EXOSC3 and EXOSC5 genes, core components of the RNA-exosome complex, which plays a major role in RNA homeostasis in the cytosol and nucleus, including quality control, degradation and processing of different RNA species (32). Both EXOSC3 and EXOSC5 interact with SARS-CoV-2 NSP8 protein which is a cofactor of the viral RNA-dependent RNA polymerase (33), suggesting these host proteins might play a role in viral RNA replication. Other recent studies have also reported direct interactions between host proteins and viral RNA and how SARS-CoV-2 infection profoundly remodels the cellular RNA-bound proteome(34).

We also identified three proteins from the toll-like receptor 4 (TLR4) signaling pathway, IRAK3, RIPK1 and NFKBIA, showing increased levels upon SARS-CoV-2 infection. TLR4 recognizes pathogen-associated molecular patterns (PAMPs) and activates innate immune systems by releasing proinflammatory cytokines via a series of events (35,36). One of the major transcriptional response to cellular stress is mediated by the heat shock response and we detected chaperones HSPA1A, HSPA9 and HSBP1 with increased staining intensity which thus suggested higher abundance upon infection, suggesting the stress response activation in infected cells as reported by previous studies (37–41).

We observed increased abundance of HS2ST1 and GLCE. From the work by Gordon and Stukalov the SARS-CoV-2 proteins NSP7 and ORF7B are shown to interact with HS2ST1 and GLCE, respectively (15,18). These proteins are known to be involved in heparin metabolism, and heparin being an anticoagulant and anti-inflammatory protein(42). Our data showing increased abundance in infected cells, supports the importance of heparin metabolism against the viral infection (43).

### Majority of host proteins show increased protein abundance and diverse spatial reorganization upon infection

Visual inspection together with computational analysis identified 10 proteins with decreased and 45 proteins with increased abundance 24 h post SARS-CoV-2 infection (**Figure 2A and Supplementary Table 1)**. Interestingly, 8 out of 10 proteins with decreased abundance were localized to mitochondria. For example the protein GRPE like 1, reported to interact with NSP10 of SARS-CoV-2 by Gordon et al (15) was downregulated significantly upon infection **(Figure 2B)** implying a potentially reduced mitochondrial function. In contrast, the 45 proteins with increased abundance were diverse in their localization. For example, IRAK3 (Interleukin-1 receptor-associated kinase), a marker for inflammation and innate immune system regulator displayed higher abundance in the plasma membrane, Golgi apparatus and ER compartment following infection (**Figure 2C**). Another target displaying significantly increased abundance was SRP72 with increased expression in both the cytosol and ER (**Figure 2A)**.

**Figure 2.**
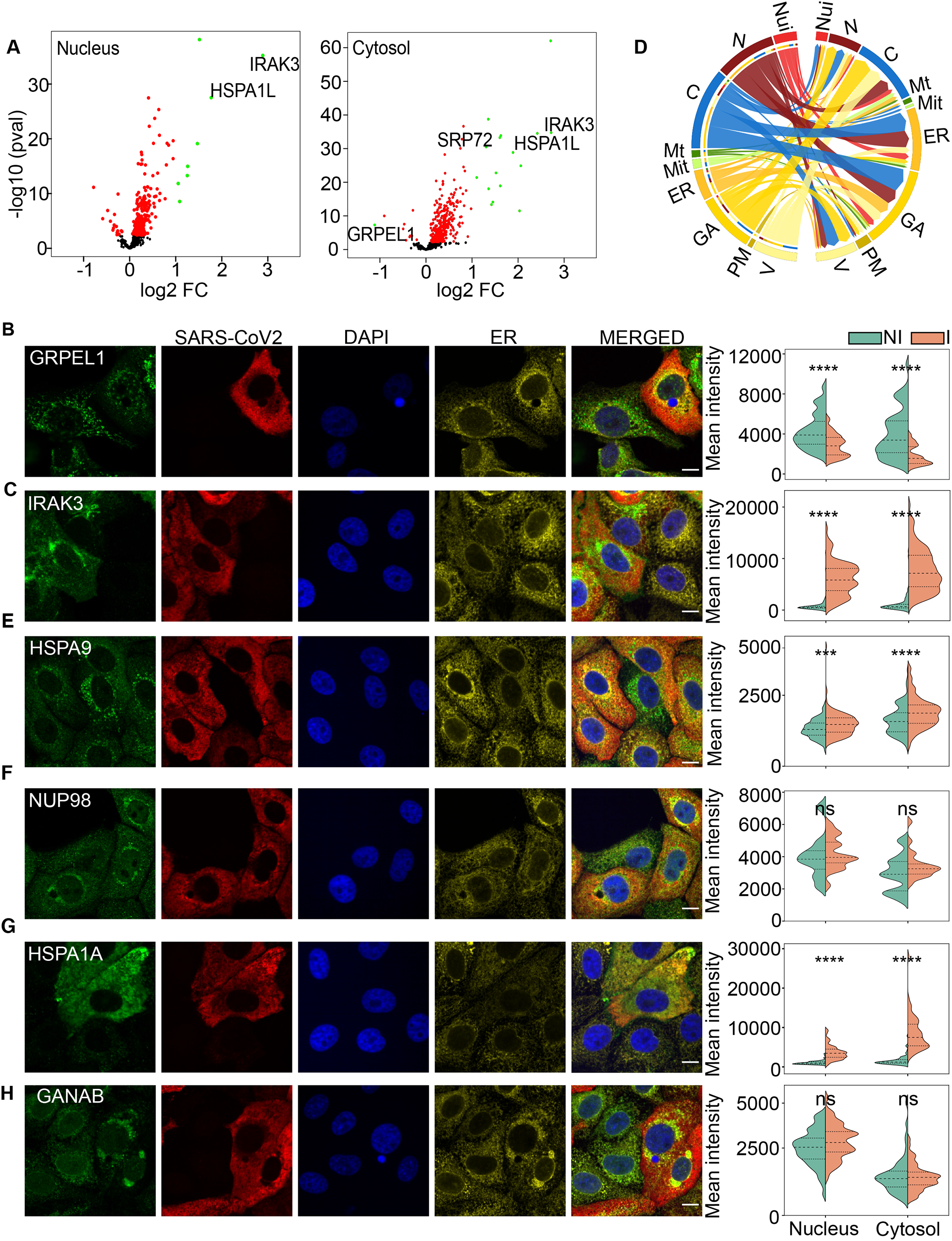
SARS-CoV-2 infection results in altered expression and translocation of many host proteins. A) Volcano plot showing overview of the host proteins with modulated expression in the nucleus and cytosol following viral infection. Proteins with significant intensity changes are shown in green. B, C) Representative images of proteins with reduced or increased intensity during SARS-CoV-2 infection, respectively. D) Circos plot showing the protein subcellular location of translocated proteins from non-infected cells (left) and infected cells (right). Nui - nucleoli, N - nucleoplasm, C - cytosol, Mt - microtubules, Mit - mitochondria, ER – endoplasmic reticulum, GA – Golgi Apparatus, PM – plasma membrane, V – vesicles. E, F) Representative images of spatially translocated proteins in the infected cells. G) Representative images of protein displaying both spatial translocation and increased intensity in infected cells. H) Representative images showing the altered ER morphology upon infection. (B, C, E-H) Host protein in green, SARS-CoV-2 NP in red, DAPI in blue, calreticulin in yellow. Scale bar represents 10 µm. The graph (right panel) shows quantified mean intensity of the host protein signal in the nucleus or cytosol for non-infected (NI) and infected (I) cells. P-value more than 0.05 = ns, less than or equal to 0.05 = *, less than or equal to 0.01 = **, less than or equal to 0.001 = *** and less than or equal to 0.0001 = ****.

Taken together our data reveal alterations in abundance for many host proteins with the majority showing higher abundance upon infection. Other proteomics studies recently published also report on a large number of significantly altered proteins upon SARS-CoV-2 infection, however with varying fraction of upregulated versus downregulated proteins depending on time point of measurement (6,14).

Further, we identified 42 proteins undergoing spatial reorganization upon infection, among which 22 proteins also showed increased abundance. A circos plot representing the reorganization of host cell proteins upon infection is shown in **Figure 2D**. Upon infection, massive reorganization of proteins occurs with a large number of proteins relocating to Golgi, ER and cytosol. For example, HSPA9 is a mitochondrial residing heat shock protein, which is partially translocating to the cytosol upon infection **(Figure 2E)**. Furthermore, the cytosolic levels vary within the population of infected cells, which could potentially be linked to viral replication cycle stage. A second example is NUP98, a protein in the nuclear pore complex, which undergoes spatial reorganization from nucleus and vesicles in non-infected cells to nucleus, vesicles and Golgi apparatus in SARS-CoV-2 infected cells (**Figure 2F)**. Also for the ER resident protein GANAB, the staining indicates a translocation to the Golgi Apparatus (**Figure 2H**). However, this may be rather a result of changed ER morphology upon infection, as the target protein stain overlaps with the ER marker used in the screen. A third example is CMPK1, a nuclear resident protein that localizes to the Golgi apparatus in non-infected cells, but also to vesicles in infected cells. Among the proteins with both spatial reorganization and altered abundance is the chaperone HSPA1A, which displays increased abundance as well as redistribution from vesicles in non-infected cells to cytosol and plasma membrane in infected cells (**Figure 2G**). Altogether, our host-response screen of 602 host cell proteins identified 97 proteins with altered protein abundance and/or subcellular distribution 24 hours post SARS-CoV-2 infection.

As mentioned above, instead of carrying all necessary elements for replication and spread, viruses hijack host cell machinery. Thus, we hypothesize the identified host cell proteins with altered spatial or expression profile to be putative targets for modulation to limit viral infection and spread.

### Drug repurposing based on host-virus interplay mapping reveals antiviral activity of rimcazole, elesclomol and crizotinib

In order to identify any available drugs designed to target the putative host proteins, the SPECS repurposing library was selected as a collection of annotated drugs. The library has been gathered based on the design criteria of the Broad Repurposing collection (44) and contains 5277 compounds including 3056 in clinical development across 600 indications and 2221 in preclinical development with varying degrees of validation. To select candidates for drug repurposing based on the cellular responses to SARS-CoV-2, drugs within the SPECS library were mapped to the 97 host proteins based on available drug annotations using the CLUE API and HIPPIE services (**Supplementary Figure 3**). Altogether, 123 drugs were identified as repurposing candidates against 21 of the 97 host proteins. The number of drugs towards proteins varied with as many as 36 drugs targeting AXL/MET, while proteins such as GANAB and CLCN7 are only targeted by one drug **(Supplementary Table 2)**.

To test the antiviral activity of the 123 drug repurposing candidates against SARS-CoV-2, our host response image based assay was transferred from 96 to 384 well plates and supplemented with a compound treatment step **(Figure 3A)**. Vero E6 cells were infected with SARS-CoV-2 in suspension and seeded onto pre-spotted compounds in duplicate for 24 hours. Infected cells treated with DMSO served as a control for infection baseline. Cells were immunostained for SARS-CoV-2 Spike protein, Calreticulin as an ER marker, and by Hoechst to identify cell nuclei. Images were subsequently acquired by high-content confocal microscopy (**Figure 3A**). Infection rate was quantified as the percentage of SARS-CoV-2 positive cells in relation to the total number of cells. For the DMSO controls, on average 80% of the cells were infected (**Figure 3B, D)**. Importantly, drug concentrations not exceeding 20% toxicity in non-infected cells were chosen for antiviral activity evaluation (data not shown). Assay quality was confirmed with Z’-factors of 0.84 and 0.90 on two assay replicates. In total, 13 out of 116 drugs significantly reduced SARS-CoV-2 infection rate to below the 3x standard deviation (SD) threshold, representing the hit compounds (hit rate 11%) while the rest of the drugs had limited activity (**Figure. 3B-D, Supplementary Figure 4A**). Crizotinib, elesclomol and rimcazole targeting AXL/MET, HSPA1A and SIGMAR1/SLC12A4, respectively, reduced SARS-CoV-2 infection nearly to the rate of non-infected cells or those treated with remdesivir; a reference antiviral compound (45,46) (**Figure 3B, D**). These host proteins responded by increased abundance in the host response screen, indicating increased expression upon SARS-CoV-2 infection (**Supplementary Table 1**). The remaining ten hit compounds showed varying activity by reducing the infection rate from 65% to 39% (**Figure 3B)**. Cell viability was reduced to 59-65 % during the treatment with crizotinib, epalrestat, ranirestat and SMI-4a, but not with the rest of the hit compounds, highlighting the need for dose-response activities to identify drug therapeutic windows **(Figure 3C, Supplementary Figure 4**). Altogether, this data presents a target-centric workflow and the identification of 13 compounds as repurposing candidates against COVID-19.

**Figure 3.**
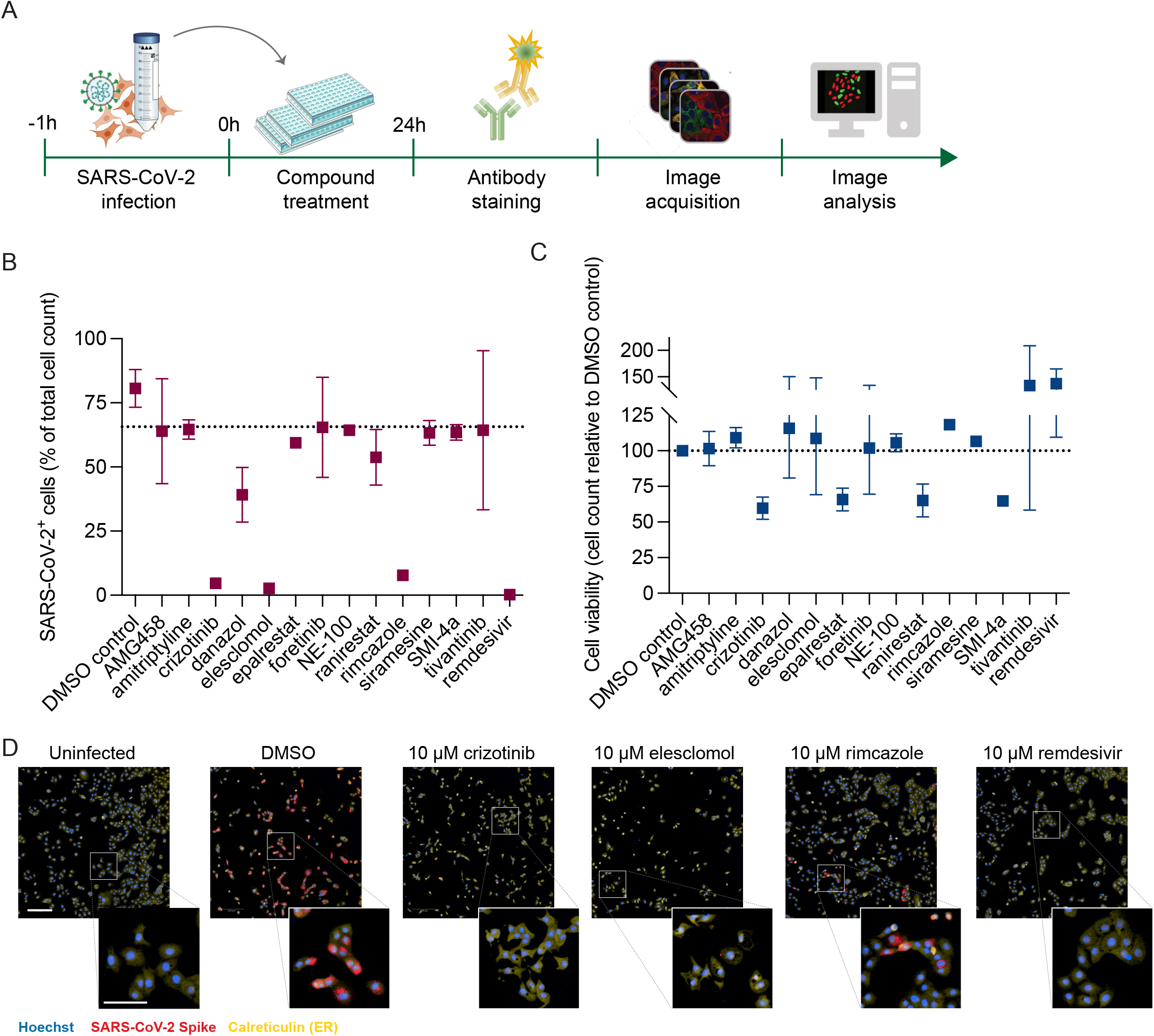
Repurposing based on host cell protein profiling reveals crizotinib, rimcazole and elesclomol antiviral activity. A) Overview workflow protocol of SARS-CoV-2 infection, compound treatment and image acquisition/analysis. (B-C) Vero E6 cells were infected with SARS-CoV-2 MOI 0.05 in suspension, seeded on 384 well-plates containing pre-dispensed compounds and incubated for 24 h. Upon fixation, cells were stained for SARS-CoV-2 Spike, Calreticulin and cell nuclei by Hoechst and analysed by high-throughput microscopy. Data is presented as mean of two technical replicates per compound from n=1 biological replicate. B) Infection rate (% total cell count) of SARS-CoV-2 infected Vero E6 cells upon drug treatment. C) Viability of Vero E6 upon drug treatment. D) Representative images of uninfected cells and SARS-CoV-2-infected Vero E6 cells treated with DMSO, crizotinib, elesclomol, rimcazole or remdesivir.

## Discussion

Diverse families of viral pathogens are well known to alter host proteome organization as part of their replication cycle (3,28) and accumulating evidence highlights the importance of protein translocations during viral infection (2). For better understanding of the host response to infection it is a golden standard to investigate virus and host proteins using bulk systematic methods such as quantitative proteomics (47,48) or sophisticated but low-throughput microscopy to unravel structure and interactions between specific viral and host cell structures or proteins at low scale (49,50). Some studies have explored the subcellular localization of the individual viral proteins using tagged versions of the proteins and immunofluorescence (47). While this adds important information about the viral proteins and complement the affinity capture methods used for studying interactions, it does not reveal changes in subcellular distribution of the host cell proteins. Most studies focusing on identifying antivirals and treatments neglect the spatial as well as single- and sub-cellular information on a systematic scale.

In this work, we leveraged spatial information at subcellular resolution during infection to build an approach for systematically shortlisting host proteins as potential antiviral targets. Mapping subcellular changes during infection enables a view inside intimately balanced homeostasis and its disturbances on an organelle, biomolecule or protein level that can guide therapeutic target or drug discovery (7). We present a novel systematic spatial profiling approach of SARS-CoV-2 infected cells to map the *in situ* landscape of host responses and subsequently demonstrate its opportunities for target-specific drug repurposing. In fact, our study feeds several potential antiviral targets to future follow-up studies as well as for drug discovery. Our data-driven approach also differs from conventional hypothesis-driven drug discovery where often a single specific target is chosen for drug screening.

By utilizing the antibody resources and expertise gathered within the HPA project and selecting antibodies specific to host proteins with previously validated interactions with SARS-CoV-2, we mapped abundance and re-localization of host responses to SARS-CoV-2 infection. Of the 602 proteins we studied, 97 changed in abundance and/or spatial re-localization, illustrating multifaceted responses in these interactions. Most proteins responded to SARS-CoV-2 with increased rather than reduced abundance which could reflect either activated cellular defense mechanisms or virus-orchestrated support to its replication machinery. We then considered the host responses as potentially druggable phenotypes, matched proteins with existing drugs from the SPECS repurposing library and identified elesclomol, rimcazole and crizotinib as drug repurposing candidates. Importantly, the approach with HPA antibodies covering most of the relevant proteome enables larger target-focused systematic screening campaigns for the discovery of new host-virus biology.

Thorough validation of target role in a disease state is undoubtedly a critical step in classical drug discovery, however studying solely the change in protein abundance and/or spatial localization can already indicate putative drug targets in the disease of interest. Utilizing the host responses to SARS-CoV-2 infection, we explored drug repurposing path by matching 21 host responses with available drug annotations in the SPECS library. Additionally, we envision the remaining 76 proteins as promising target validation and drug discovery starting points. For instance, Polypyrimidine tract-binding protein 2 (PTBP2), an RNA binding protein involved in pre-mRNA splicing, has been described as a potential novel drug target by Gordon *et al* study (15). The drug repurposing proof-of-concept identified three categories of drug-target links being relevant for coronavirus infection as well as infectious diseases generally. We anticipate how crizotinib, elesclomol and rimcazole could exhibit activity via three possible mechanisms: inhibition of their annotated target(s), effects on unknown proteins or by poly-pharmacological activity.

As a multi-target tyrosine kinase inhibitor, crizotinib is known to target AXL, MET, ALK and RON (the latter two not explored in this study) used in the treatment of ALK-positive metastatic non-small cell lung cancers (51). In SARS-CoV-2 infection, AXL interaction with the Spike protein has shown to be crucial for virus entry into cells and supportive of viral infection in primary lung epithelium (52). Another AXL inhibitor, bemcentinib, has the potential to increase the survival of COVID-19 patients compared to standard of care as presented in Phase II trial by BerGenBio (Clinical Trial: NCT04890509), supporting AXL-mediated antiviral activity of crizotinib. Still, crizotinib is only one out of seven AXL inhibitors having an antiviral activity in our study and thereby seems to exhibit diverse activity on distinct target(s) in their primary disease indication and COVID-19 patients.

Rimcazole is a carbazole derivative acting partially as a SIGMAR1 antagonist with additional affinity for dopamine transporters which was discontinued as an anti-schizophrenia drug in early 1980s due to lack of efficacy(53). In fact, other SIGMAR1 inhibitors, but not rimcazole, have been previously identified as antivirally active in *in vitro* repurposing screens against HCV, Ebola virus (EBOV) and coronaviruses, including SARS-CoV-2 (54–58). However, concerns were recently raised around SIGMAR1 inhibitors when induction of phospholipidosis was described to underlie the antiviral activity of most of the proposed repurposing candidates (59). Similarly to crizotinib, rimcazole is the only one out of 22 SIGMAR1 inhibitors that presented antiviral activity.

Elesclomol is mainly known as a pre-clinical anti-cancer drug candidate inducing oxidative stress in mitochondria (60), triggering apoptosis in cancer cells and activation of heat shock proteins and signaling pathway (61). Increased abundance of HSPA1A gene encoding for the major cytosolic HSP70 protein upon SARS-CoV-2 infection, as well as elesclomol antiviral activity indicates HSP70 putative role in the virus infection cycle. We speculate the elevated expression levels of heat shock proteins to be part of the cellular defense mechanism, aiding in targeting viral proteins for degradation, rather than assisting in folding of the viral proteins.

A limitation of this study is that results are based on data from the non-human cell line Vero E6, originating from the African green monkey. This cell line is known for its dampened innate immune response and permissiveness to SARS-CoV-2, which makes it a feasible but a limiting virology model (62). When comparing subcellular localization of the hits from the host-response screen between non-infected Vero E6 cells and the immunofluorescence data on human cell lines as previously generated within the HPA, 80% of the patterns overlap between the species (data publicly available at *www.proteinatlas.org*). Looking at staining similarities across all proteins (n=546) stained in Vero E6 in addition to human cell lines within the HPA, 75% show overlap in subcellular localization. Due to the inter-species similarities, we speculate that host proteome landscape responses to SARS-CoV-2 infection in a partially similar manner in human cell lines.

In this work we performed targeted phenotyping of disease relevant proteins as a funnel to guide target selection for drug repurposing or discovery. Further, we suggest that “targeted phenotyping” can be used to prioritize host targets for novel drugs, in this case for the treatment of COVID-19.

Our approach can be applied as a stand-alone filter or as an integrated layer in multi-omics study for the selection of relevant host responses in infectious diseases. Given that the approach is easily scalable and transferable for infectious agents or other diseases beyond SARS-CoV-2, we anticipate that the spatial dimension will support fitting a crucial piece in the puzzle of various diseases.

## Materials and Methods

### Cell culture and virus infection

Vero E6 cell line was grown at 37°C in a 5% CO_2_ condition in Dulbecco’ modified Eagle’s medium (DMEM) containing 10% FBS (Gibco). For the host protein response screen 10,000 cells/well were seeded on 96 well microplates (Perkin Elmer, CellCarrier-96 Ultra Microplates, tissue culture treated, product number 6055302) and incubated for 20 - 24 h. The cells were washed twice with PBS and infected with SARS-CoV-2 at MOI 0.001 for 1 h at 37 °C. The SARS-CoV-2 strain (CoV-2/human/SWE/01/2020) was obtained from the Public Health Agency of Sweden.

The virus containing media was removed and washed twice with PBS followed by replenishing with fresh media containing 2% FBS and incubated for 24 h. After washing twice with PBS the cells were fixed with 4% paraformaldehyde (PFA) (Sigma Aldrich, Darmstadt, Germany) for 1 h, followed by washing with PBS. The 96-well assay was adopted to a 384-well format suitable for drug screening by introducing a drug treatment step and by changing the infection procedure. For the experiments with drug treatments, cells were detached using TryplE (Gibco, USA) and resuspended in fresh media without FBS and infected with SARS-CoV-2 at MOI 0.05 for 1 h at 37 °C. The virus containing media was removed and the cells were washed twice with PBS followed by replenishing with fresh media containing 10% FBS. 30 µL containing 2,500 cells/well were seeded on 384 well-plates (Perkin Elmer, PhenoPlate 384-well microplates) containing pre-dispensed compounds dissolved in DMSO.

### Antibody selection

Validated antibodies from the HPA project were blasted against the *Chlorocebus sabaeus* sequence from Ensembl. Proteins with more than 60% identity across the whole length of the antigen sequences used to generate the HPA antibody were selected.

### Immunofluorescence staining

#### Host response screen

Cells were fixed, permeabilized and stained as previously described (19). Briefy, fixed cells were washed with PBS, permeabilized using 0.1% Triton X-100 (Sigma Aldrich) in PBS for a total of 15 min with new Triton solution added every 5 minutes (3 × 5 minutes). very and washed again with PBS by using a Tecan Evo Freedom pipetting workstation. Cells were then incubated in primary antibodies overnight in PBS containing 4% FBS. Rabbit polyclonal primary antibodies generated at the HPA were diluted in concentration of 2 µg/ml. The commercial antibody towards the Sigma-1 R (MAB1076, R&D systems) was diluted to 3 µg/ml, chicken Calreticulin (Abcam, ab2908) to 1 µg/ml and mouse SARS-CoV-2 Nucleoprotein monoclonal antibody (MA1-7404, Thermofisher) to 10 µg/ml. Cells were then washed 4X with PBS and incubated with secondary antibodies for 90 min in dark at RT. Secondary antibodies goat anti□mouse Alexa Fluor 555 (A21424), goat anti□rabbit Alexa Fluor 488 (A11034), and goat anti□chicken Alexa Fluor 647 (A21449), were diluted to 2.5 µg/ml in PBS containing 4% FBS. Following, cells were incubated with 0.2 µg/ml DAPI for 10 min and washed 4X with PBS. Plates were sealed and stored at +4°C until image acquisition.

#### Drug repurposing screen

Fixed cells were washed with PBS, permeabilized using 0.1% Triton X-100 (T8787, Sigma-Aldrich) in PBS for 15 min, washed with PBS and blocked in 4% BSA (A7030, Sigma-Aldrich) for 1 h at RT. Cells were then incubated with primary antibodies overnight at 4°C. Primary antibodies generated at the HPA were diluted to 2 µg/ml, commercial Sigma-1 R antibody to 3 µg/ml (MAB1076, R&D systems) chicken Calreticulin (Abcam, ab2908) to 1 µg/ml and mouse SARS-CoV-2 Spike monoclonal antibody (GTX632604 GeneTex) to 1 µg/ml. Upon primary antibody incubation, cells were washed 6x with PBS and incubated with secondary antibodies for 90 min in dark at RT. Secondary antibodies goat anti□mouse Alexa Fluor 555 (A21424), goat anti□rabbit Alexa Fluor 488 (A11034), and goat anti□chicken Alexa Fluor 647 (A21449), were diluted to 2.5 µg/ml in PBS containing 4% BSA. Following, cells were incubated with 1 µg/mL Hoechst 33342 for 10 min and washed 4X with PBS. Plates were sealed and stored at +4°C until image acquisition.

### Image acquisition

Immunostained cells were imaged in PBS using a laser-based Opera Phenix high-content microscope (PerkinElmer) in confocal mode with a 63X water objective. Nine to twelve fields of view were imaged per well (corresponding to a few hundred to thousand individual cells), at three different z-planes to ensure proper focus throughout the automatic image acquisition across the plates. Raw 16-bit images were exported as TIFF files and the z-planes were combined to create max projections prior to analysis. All raw data images including all individual z-planes are available on Figshare (22) (https://scilifelab.figshare.com) under the following doi: 10.17044/scilifelab.14315777.

For the drug repurposing screen images were additionally acquired with a 10X air objective and four fields of view per well for inclusion of the entire cell population per well.

### Cell segmentation and Image quantification host protein interaction screen

The acquired images were transferred to an application built in the ImJoy platform (21) (a server-based web application) for manual annotation of the subcellular locations of each protein under investigation.

For quantification of relative protein expression in infected and non-infected cells, images were segmented to identify individual cells, as well as the regions of the nucleus and the cytoplasm. We used a DPNUnet model trained with manually segmented HPA images) to generate binary cell masks for each image. The segmentation masks were generated for the nucleus by using the DAPI channel and the whole cell by using the ER channel as input. The cytoplasm was defined by subtracting the nuclear region from the whole cell. Intensities for the target protein, ER and SARS-CoV-2 channels were then quantified separately for these regions, both as mean and integrated values. To define infected and non-infected cells, we calculated the mean pixel values for each cell by using the third quartile (51% to 75% highest values (above the median) pixel values for the virus channel in the region of the cytoplasm. Based on the value a threshold is set to define whether cells are infected or not. The segmentation panel marks the infected and non-infected cells differently to verify the segmentation model and enables further fine tuning of the threshold value if needed. For each well, protein staining intensity was quantified for infected and non-infected cell populations and the fold change between the populations was calculated. A t-test was done to calculate the statistical significance between the infected and non-infected populations and the p-value was adjusted for false discovery rate with Benjamini Hochberg. Proteins with adjusted *p*-value < 0.01 and log2 of fold change > 1 were considered upregulated. Proteins with adjusted *p*-value < 0.01 and log2 of fold change < 1 were considered downregulated. Violin plots were generated and the overall distribution of fold change for the analysed proteins were displayed in a volcano plot.

During manual annotation of protein subcellular localization, annotators assigned the population of infected and non-infected cells to one or multiple subcellular organelles, which included nucleoplasm, nuclear membrane, nucleoli, nucleoli fibrillar center, nuclear speckles, nuclear bodies, kinetochore, mitotic chromosome, endoplasmic reticulum, Golgi apparatus, vesicles, peroxisomes, endosomes, lysosomes, intermediate filaments, actin filaments, focal adhesion sites, microtubules, microtubule ends, cytokinetic bridge, midbody, midbody ring, cleavage furrow, mitotic spindle, primary cilia, centriolar satellites, centrosome, lipid droplets, plasma membrane, aggresome, cytosol, mitochondria, cytosol, cytosolic bodies and rods and rings. The proteins were categorized as spatial hits if the subcellular locations were annotated differently between non-infected and infected cell populations.

### Generation of virus and host protein interaction network

Viral bait - host protein and host protein-protein interaction network was visualized with Cytoscape (63). SARS-Cov-2 Viral bait - host protein interactions were derived from two recently published papers by Gordon et al. and Stukalov et al (15,18). with a significant threshold of 0.05. Human protein - protein interactions were derived from String database (64). Protein complexes and biological process grouping were derived from CyCommunityDetection (23). In specific, communities were detected with Louvain, Infomap and HiDeF. Functional enrichment were determined by the robust functional groups found by Enrichr (65) out of Gene Ontology databases (GO_Biological_Process_2018, GO_Cellular_Component_2018, GO_Molecular_Function_2018), KEGG database (KEGG_2019_Human), Reactome database (66), WikiPathways and Human Phenotype Ontology.

### Quantification of SARS-CoV-2 infection upon compound treatment

Image analysis was performed using Harmony software (PerkinElmer). Cell nuclei were identified using the Hoechst 33242 channel, through application of the “Find nuclei” algorithm. Cell boundaries were identified using the Alexa 647 channel detecting Calreticulin signal through the application of the “Find cytoplasm” algorithm. To distinguish infected and non-infected cells, average intensity of the Alexa 555 channel detecting SARS-CoV-2 Spike protein was calculated for the perinuclear space by a 60% increased area around the nuclei with applied threshold. After image analysis, per-well data was used to calculate cell viability and infection rate. Cell viability was calculated as percentage from the total number of cells in DMSO-treated control and infection rate was calculated as SARS-CoV-2^+^ cell percentage from the total number of cells in corresponding treatment. Data was plotted using Graphpad Prism software.

### Drug library annotation

The SPECS repurposing library was obtained from Chemical Biology Consortium Sweden. The 5291 available drugs were annotated using the CLUE API service (https://clue.io/developer-resources#apisection). In detail, for 5114 compounds with an available pert_iname, we first retrieved compound information, mechanism of action and targets from the CLUE database. For 3674 compounds, a unique set of 1895 gene symbols were reported. To visualize the connections between drugs and protein targets, a Protein-Protein Interaction (PPI) network was obtained from HIPPIE (http://cbdm-01.zdv.uni-mainz.de/~mschaefer/hippie/index.php, v2.2). Then, a subnetwork composed of the 97 proteins and their adjacent edges was considered. Interactions with compounds were then added (**Supplementary Figure 3**) considering the CLUE target gene names.

### Compound handling for drug repurposing screen

Assay ready plates with pre-dispensed compounds were prepared in black 384-well imaging plates (Perkin Elmer, PhenoPlate 384-well microplates, tissue culture treated, product number 6057302) using acoustic dispensing (Echo 550, Labcyte). For investigational compounds from the SPECS library, the assay ready plates contained either 2.5 nL or 30 nL of compound stock solutions in DMSO (10 mM or 0.1 mM), corresponding to non-toxic doses chosen based on toxicity evaluation in non-infected cells (data not shown). Reference compound remdesivir was spotted in 30 nL of 10 mM stock solution in DMSO as an antiviral reference compound. DMSO was used as negative control and spotted in 30 nL. Spotting volumes, stock concentrations and final assay concentrations for all drugs are summarized in Supplementary Table 2. The plates were then heat-sealed using a peelable aluminum seal (Eppendorf, 0030127790) with a thermal microplate sealer (PlateLoc, Agilent) and then stored at -20 °C until use. On the day of the experiment the plates were allowed to thaw for 30 minutes prior to removal of the seal.

## Figure legends

**Supplementary Figure 1.**
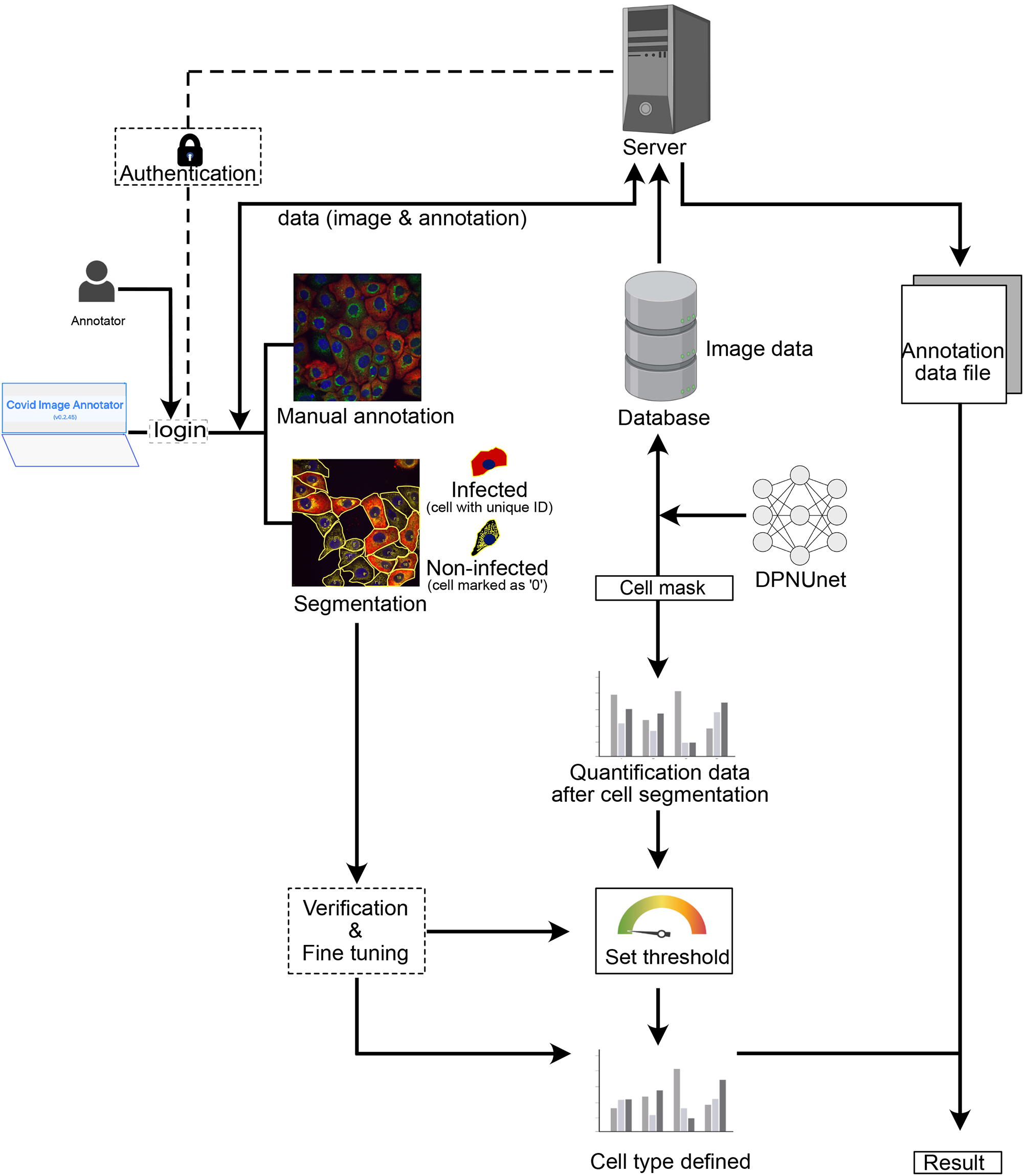
Overview of the image analysis pipeline starting from imaging to computational analysis and generation of final results.

**Supplementary Figure 2.**
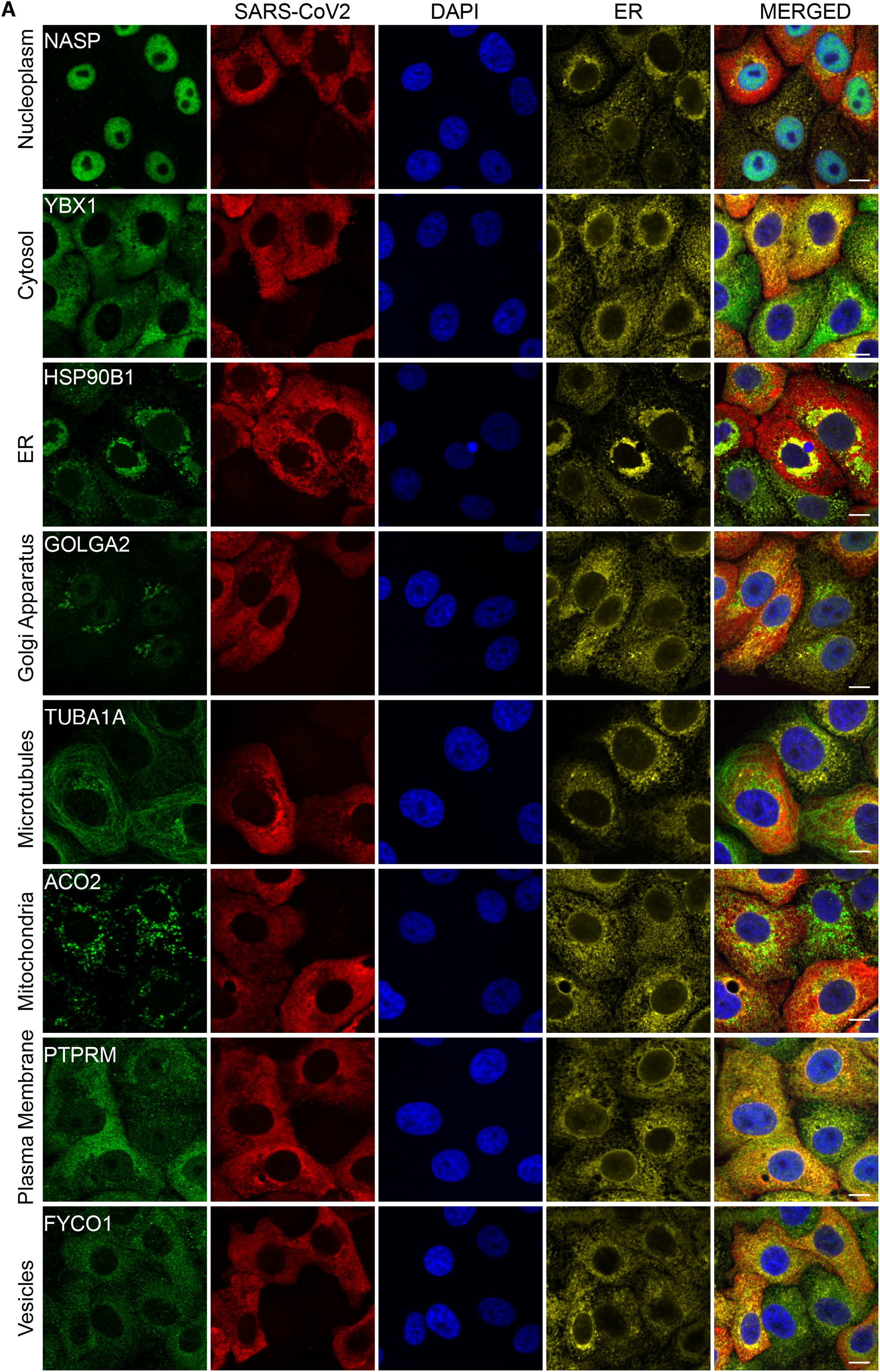
Examples for organelle markers namely nucleoplasm, cytosol, endoplasmic reticulum, Golgi apparatus, microtubules, mitochondria, plasma membrane, vesicles. Scale bar represents 10 µm.

**Supplementary Figure 3.**
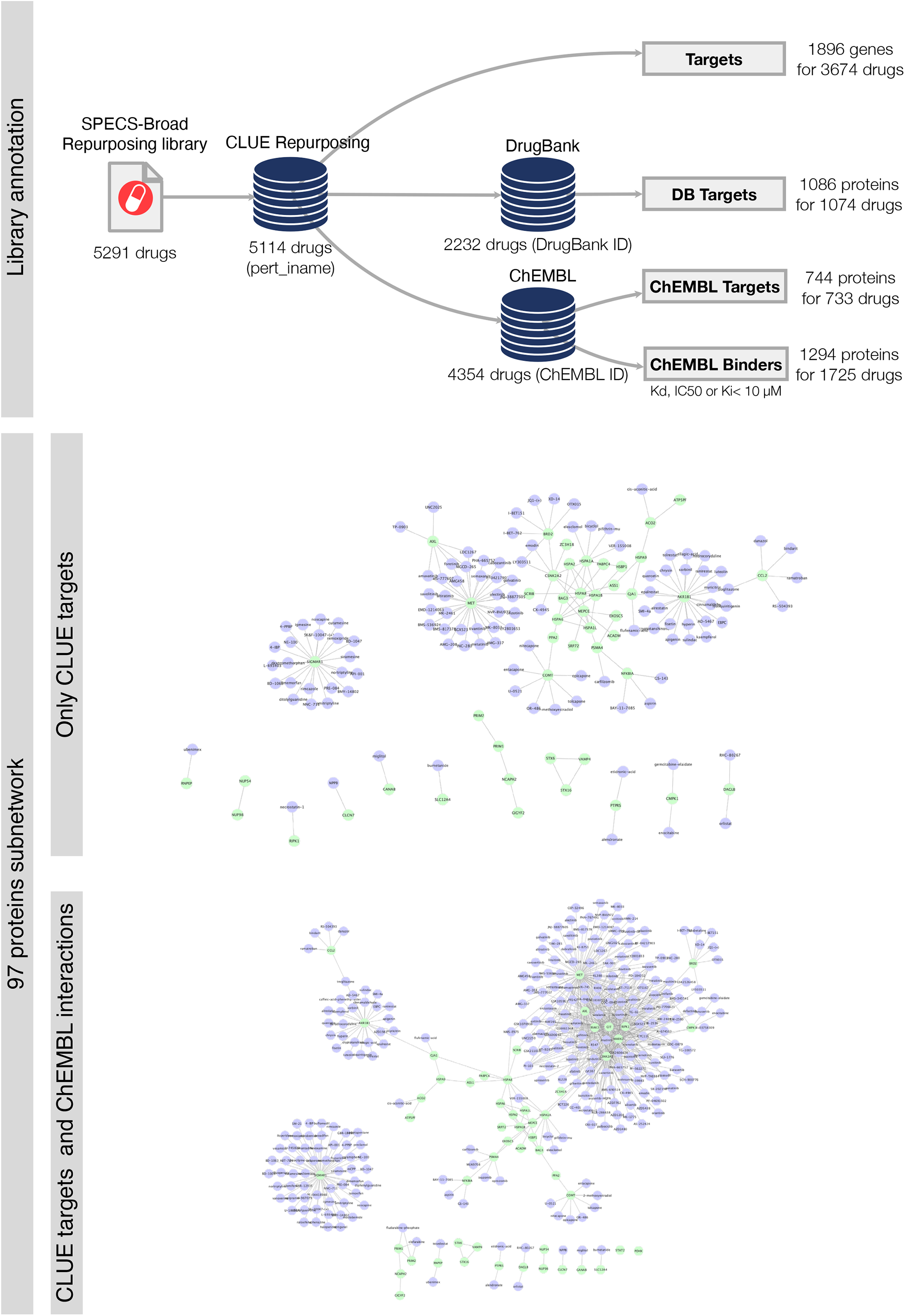
A) Workflow of matching host targets with existing drugs in the SPECS repurposing library (see details in Materials and Methods and Supplementary Table 2). In the network visualization single unconnected proteins or proteins not available in the reference PPI are not shown. For following drug repurposing screen, only annotations retrieved from the Clue API database were used.

**Supplementary Figure 4.**
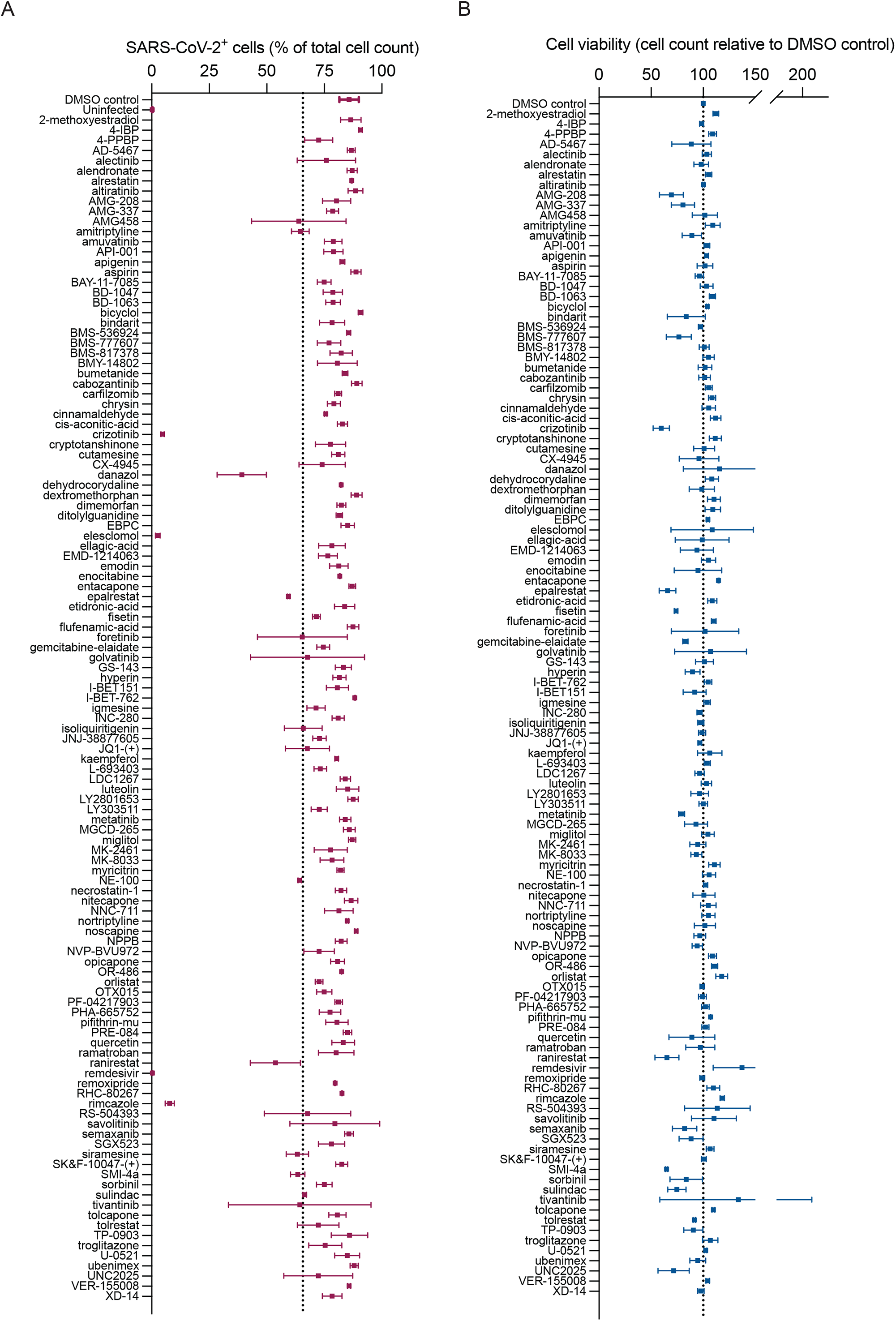
Antiviral activity and cellular viability response of Vero E6 cells treated with selected compounds in the SPECS library. (A-B) Vero E6 cells were infected with SARS-CoV-2 MOI 0.05 in suspension, seeded on 384 well-plates containing pre-dispensed compounds and incubated for 24 h. Upon fixation, cells were stained for SARS-CoV-2 Spike, Calreticulin and cell nuclei by Hoechst and analysed by high-throughput microscopy. Data is presented as the mean of two technical replicates per compound from n=1 biological replicate. A) Infection rate (% of total cell count) of SARS-CoV-2 infected Vero E6 cells upon drug treatment. B) Viability of Vero E6 upon drug treatment.

**Supplementary Table 1. Intensity and spatial information summary of all evaluated proteins**. Summary of all proteins and antibodies used in the study with protein mean intensity in the nucleus, cytosol, and cell areas (Sheet 1). Localization details of all proteins in infected and non-infected cells (Sheet 2).

**Supplementary Table 2. Summary of gene targets, corresponding drugs and concentrations used in the drug repurposing screen**. Drug target annotations based on CLUE (Sheet 1), Drug concentrations (Sheet 2).

## Acknowledgements

In vitro work for the repurposing screen with active SARS-CoV-2 was performed at the Biomedicum BSL-3 Core Facility, Karolinska Institutet. The compound library and plating thereof was provided by the Chemical Biology Consortium Sweden (CBCS). We also acknowledge the Human Protein Atlas, funded by Knut and Alice Wallenberg foundation for the use of all primary antibodies in the screen. We acknowledge the Spatial Proteomics Facility, funded by SciLifelab for optimizing and performing the host protein identification screen and Knut and Alice Wallenberg for funding of the particular study (KAW 2020-0182). This work has also been supported by EPIC-XS, project number 823839, funded by the Horizon 2020 programme of the European Union.

